# Immune-active microenvironment in small cell carcinoma of the ovary, hypercalcemic type: rationale for immune checkpoint blockade

**DOI:** 10.1101/197970

**Authors:** Petar Jelinic, Jacob Ricca, Elke Van Oudenhove, Narciso Olvera, Taha Merghoub, Douglas A. Levine, Dmitriy Zamarin

## Abstract

Small cell carcinoma of the ovary, hypercalcemic type (SCCOHT) is a highly aggressive monogenic cancer driven by *SMARCA4* mutations. Here, we report responses to anti-PD1 immunotherapy in four patients and characterize the immune landscape of SCCOHT tumors using quantitative immunofluorescence and gene expression profiling. Unexpectedly for a low mutation burden cancer, the majority of the tumors demonstrated PD-L1 expression with strong associated T cell infiltration. PD-L1 expression was detected in both tumor and stromal cells, with macrophages being the most abundant PD-L1 positive cells in some tumors. Transcriptional profiling revealed increased expression of genes related to Th1 and cytotoxic cell function in PD-L1 high tumors, suggesting that PD-L1 acts as a pathway of adaptive immune resistance in SCCOHT. These findings suggest that although SCCOHT are low-mutational-burden tumors, their immunogenic microenvironment that resembles the landscape of tumors that respond well to treatment with PD-1/PD-L1 blockade.

---

Small cell carcinoma of the ovary, hypercalcemic type (SCCOHT) is a rare and very aggressive malignancy that occurs mostly in young women [1]. We and others previously described SCCOHT as a *SMARCA4*-mutated monogenic disease and unraveled other important molecular features [2-6]. Despite these advances in understanding the biology of SCCOHT, there are few effective treatment options and survival rates remain poor.

Immunotherapy utilizing antibodies against the immune checkpoint inhibitor programmed death 1 (PD-1) has emerged as a promising treatment option against many human cancers [7-12]. High tumor mutational load, presence of tumor infiltrating lymphocytes (TILs), and PD-L1 expression in the tumor microenvironment appear to enrich for patients responding to PD-1/PD-L1 blockade [13-15]. Studies in gynecologic malignancies, including endometrial cancers with microsatellite instability (MSI) and BRCA-mutated ovarian cancer demonstrate that an increase in mutational burden appears to be associated with an increase in TILs and tumor PD-L1 expression, creating a rationale for immunotherapy with PD-1/PD-L1 blockade [16-18].

SCCOHT is a monogenic disease and as such is also a non-hypermutated cancer [2-6], which may otherwise dampen enthusiasm for immunotherapy. The activity of PD-1/PD-L1 blockade in these tumors has not been published, yet several patients with SCCOHT have received anti-PD1 immunotherapy and early outcomes suggest benefit from these drugs. We collected case histories from four patients diagnosed at ages 18 to 29 years. Each patient was found to have a large ovarian tumor and underwent surgery that resulted in complete tumor resection. Three of the four patients received adjuvant treatment with multi-agent cytotoxic chemotherapy. All patients recurred after disease-free interval of one to three years. Each patient received additional therapy with either cytotoxic or targeted agents followed by radiotherapy and then began treatment with anti-PD1 immunotherapy. One patient has a sustained partial response for 6 months and the three other patients remain disease-free for 1.5 years or more. More detailed case histories can be found in the Supplementary data.

These anecdotal findings prompted us to study the tumor microenvironment to examine the rationale for immunotherapeutic approaches in SCCOHT. Using microscopy with immunohistochemistry (IHC) and immunofluorescence (IF) analysis, we assessed tumor expression of PD-L1, CD3 (T-cells) and CD68 (macrophages) in eleven SCCOHT cases. All but one of the patients demonstrated either a deleterious mutation in *SMARCA4*, or loss of SMARCA4 expression by IHC (Supplementary Table 1, available online) without any additional genetic alterations detected on a targeted mutation panel encompassing more than 400 genes, highlighting the likely low overall mutational burden of these tumors. Case 107 had an in-frame 4 amino acid deletion in SMARCA4 within a non-essential region and SMARCA4 IHC was equivocal due to poor quality material.

Unexpectedly for a cancer with few somatic mutations, PD-L1 expression and prominent T cell infiltration were detected in most tumors (Figure 1, A, B and C). The majority of the tumors also demonstrated infiltration by CD68+ macrophages (Figure 1D). Similar to reports in other solid tumors, PD-L1 expression was detected in both tumor cells and stromal cells, and in two cases tumor associated macrophages were the most abundant PD-L1 positive cells (Figure 1, A and E; Supplementary Figure 1, available online).

**Figure 1.**
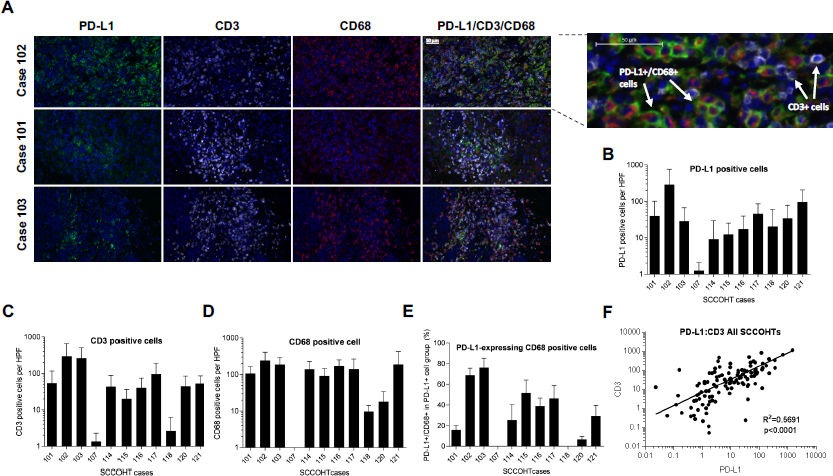
SCCOHTs exhibit immune-active tumor microenvironment. **A)** Immunohistochemistry triple-staining with PD-L1, CD3 and CD68 antibodies of three representative SCCOHT cases. Enlarged outset demonstrates merged triple-stain, showing overlap of PD-L1- and CD68-positive cells but not CD3-positive cells. **B)** PD-L1-positive, **C)** CD3-positive and **D)** CD68 positive cell counts per high power field (HPF). The y-axis indicates number of marker-positive cells and the x-axis indicates SCCOHT cases. Error bars represent one SD per 10 HPFs. **E)** Percent of CD68+ cells out of all PD-L1-expressing cells. The y-axis indicates percentage of PD-L1-expressing macrophages and the x-axis indicates SCCOHT cases. Error bars represent one SD per 10 HPFs. **F)** Correlation of PD-L1 and CD3 expression in all SCCOHT cases. Each dot represents an individual HPF.

There was a strong association between T cell infiltration and PD-L1 expression across all patients (Figure 1F), and on per field basis in each patient, apart from those with the lowest PD-L1 positivity (Supplementary Figure 2A, available online). There was poor association between PD-L1 expression and tumor macrophage infiltration (Supplementary Figure 2B, available online).

**Figure 2.**
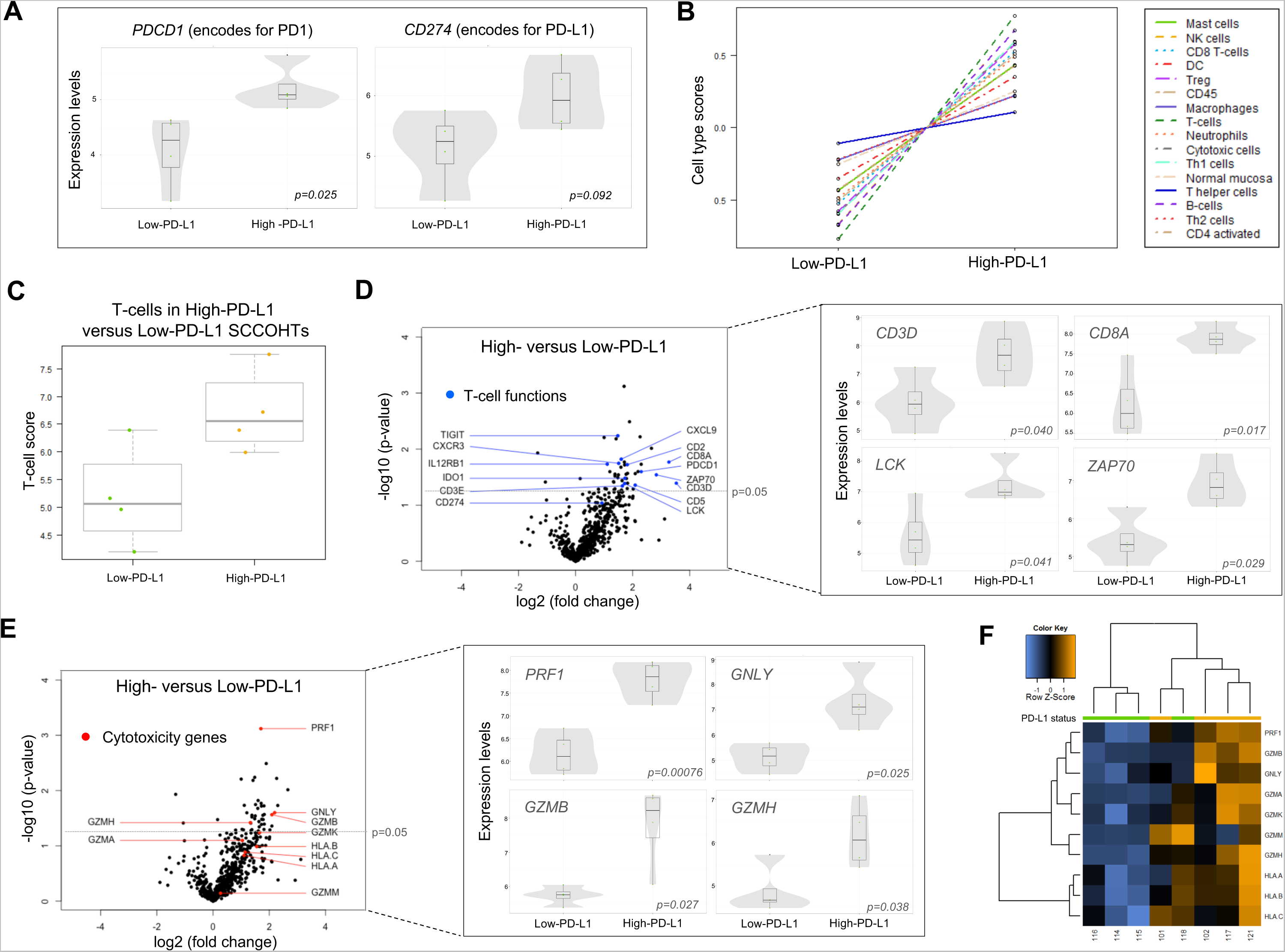
PD-L1 expression is associated with T cell-related gene expression in SCCOHTs **A)** Normalized gene expression of *PDCD1* (encoding for PD1) and *CD274* (encoding for PD-L1) in the High-PD-L1 versus Low-PD-L1 cases. Box plots superimposed on violin plots in figures A), D) and E) represent differential gene expression. Boxplot center lines represent tumor medians, box limits are the interquartile range from 25% and 75%, whiskers represent the extent of tumors out to 1.5 times the inter-quartile range. The grey shading of the violin plots represents the estimated distribution of the expression values. Dots represent individual cases. **B)** Relative abundance of specific cell types in High-PD-L1 versus Low-PD-L1 cases. Cell type scores are mean centered based on deconvolution of cell-type specific gene sets. **C)** T-cell scores in the High-PD-L1 versus Low-PD-L1 group. **D) and E)** Volcano plots of differentially expressed genes showing the T-cell-related genes (blue dots) and cytotoxicity genes (red dots), respectively. Expression of representative genes are shown to the right of each plot. Statistically significant genes are shown above the dashed line (p=0.05). **F)** Unsupervised clustering of cytotoxicity-related and MHC class I genes for High-PD-L1 and Low-PD-L1 cases. PD-L1 group is indicated by the color bar at the top of the figure (green: Low-PD-L1, orange: High-PD-L1).

The association between PD-L1 expression and T cell infiltration has been reported previously in other cancers and suggests that PD-L1 expression in SCCOHT is likely not intrinsic, but rather represents a mechanism of adaptive immune resistance to TILs [13, 19]. To characterize the tumor inflammatory responses and their association with PD-L1 expression in more detail, we performed gene expression profiling using NanoString’s nCounter PanCancer Immunoprofiling panel assessing more than 700 immune-related genes (Figure 2). We stratified the SCCOHT cases into two groups; the four highest (101, 102, 117 and 121) versus four lowest (114, 115, 116 and 118) PD-L1-expressing cases. Out of 36 differentially expressed genes (p < 0.05), 34 were elevated in the High versus Low group, the majority of which were T-cell-specific or cytotoxicity-related genes (Supplementary Table 2, available online.). The expression of *PDCD1* (encoding for PD-1) was higher in the High-PD-L1 group (Figure 2A), which has also been previously reported to be associated with response to PD-1 blockade [13]. *CD274* (encoding for PD-L1) also had higher expression in the High-PD-L1 group, however, this did not reach a statistical significance, possibly due to heterogeneity of the PD-L1-positive cells or post-transcriptional regulation of PD-L1 expression. Gene-expression based cell type deconvolution demonstrated that all immune cell types were more common in the High versus Low group, further confirming correlation between PD-L1 positivity and enhanced immune response (Figure 2B). Cell types known to be involved in anti-tumor activity (T-cells, Th1, CD8 and cytotoxic cells) were particularly elevated (Figure 2, C and D; Supplementary Figure 2C, available online). Two genes encoding for CD3, *CD3E* and *CD3D*, were elevated in the High PD-L1 group, supporting our observations from the IHC experiments that TILs and PD-L1-positive cells co-exist in SCCOHTs (Figure 2D). Cytolytic and antigen-presenting genes were also noted to be elevated in the High-PD-L1 group (Figure 2, E and F). Among cytotoxic genes, a 2-4 fold increase in perforin 1 (*PRF1*), granzyme B (*GZMB*), granulysin (*GNLY*) and granzyme H (*GZMH*) were the most prominent and statistically significant (Figure 2E), highlighting the association between PD-L1 expression and cytotoxic cell activity.

Finally, the KEGG pathway analysis revealed increased activity in the T-cell receptor signaling pathway, further strengthening our observation that TILs are engaged in active immune response in SCCOHTs (Supplementary Figure 3, available online).

These data suggest that SCCOHTs are immunogenic tumors and exhibit significant levels of T cell infiltration and PD-L1 expression. These findings are somewhat unexpected, given the low mutational burden (with likely resulting low neoantigen burden) in these tumors. Notably, in our analyses, case 107 was an outlier with both the lowest PD-L1 expression and the lowest number of TILs and macrophages. Since this case may have potentially retained SMARCA4 functionality, it follows that the transcriptional program regulated by SMARCA4 may influence tumor immunogenicity, leading to TIL infiltration and PD-L1 upregulation.

Overall, these findings contribute to understanding the immune landscape of SCCOHT tumors. The SCCOHT tumor microenvironment resembles other immunogenic tumors that respond well to anti-PD-1 immunotherapies, generating a strong rationale for evaluation of these agents in SCCOHT patients. These findings further highlight the inconsistencies between mutational burden and immunogenicity, calling for identification of additional biomarkers to assess tumor recognition by the immune system.

## Notes

We appreciate the support of all the patients who shared biologic specimens and clinical information for this report. We are thankful to Yanyun Li for help with pathology and quantification.

## Supplementary Methods

### Patients and tumor samples

In this study, we included 11 SCCOHT cases (Supplementary Table 1, available online), of which 5 (cases 101, 102, 103, 107 and 114) have been previously reviewed and analyzed for *SMARCA4* mutations [1, 2]. Using previously described guidelines [1], specialty gynecologic pathologists reviewed new cases — 115, 116, 117, 118, 120 and 121 — to confirm the diagnosis of SCCOHT. Clinical data collection was limited to only age due to the Health Insurance Portability and Accountability Act regulations. Specific case details were collected directly from involved patients and deemed to be IRB-exempt.

### MSK-IMPACT Assay and Sequencing

For the purposes of the MSK-IMPACT assay, DNA was extracted from formalin-fixed paraffin-embedded (FFPE) tumors with at least 50% tumor cell nuclei according to standard protocols (DNeasy Blood&Tissue kit; Qiagen #69506). The assay was performed as previously described [2]. Briefly, barcoded sequence libraries were prepared using 250 ng of genomic DNA (Kapa Biosystems, Wilmington, MA, USA) and combined into a single equimolar pool. The captured pool was subsequently sequenced on an Illumina HiSeq 2500 sequencer (Illumina, San Diego, CA, USA) as paired-end 100-base pair reads, producing at least 250-fold coverage per tumor. A panel of 341 known oncogenes and tumor-suppressor genes frequently altered in cancer were sequenced.

### Immunostaining

All assays were performed on formalin-fixed paraffin-embedded slides. Immunohistochemistry staining for SMARCA4 was performed as previously described [1]. Immunofluorescent staining for human CD3 (rabbit polyclonal, DAKO #A0452; 1.2 µg/ml), CD68 (mouse clone KP1, DAKO #M0814; 0.02 µg/ml), and PD-L1 (rabbit clone E1L3N, Cell Signaling Technology #13684; 1.67 µg/ml) were performed with an automated Leica Bond RX processor using a published protocol [3]. Slides were digitized using Pannoramic Flash scanner (3DHistech). Quantification of CD3, CD68, and PD-L1 positive cells was performed by selection of 10 random high power fields (HPFs) in each sample. The random HPFs were converted to 8-bit images and the areas of each signal (DAPI, A488, A594 or A568, and overlapping A488 and A594 or A568) were calculated for each field using ImageJ software. Multiplier constants specific for each stain (CD3, CD68, PD-L1 and double positive CD68 and PD-L1) were used to correct for differences in each signal area compared to that of DAPI for a single cell.

These were calculated by dividing the total signal area by the average DAPI area per cell. The estimated cell counts were validated by manual counts by two independent evaluators, including one pathologist, in at least 20 random HPFs from all samples. Percent positive signal was calculated by multiplying the total signal area by the appropriate multiplier constant and then dividing by the total DAPI area within each field. Estimated positive cell counts were calculated by multiplying the total signal area by the appropriate multiplier constant and then dividing by the average DAPI area per cell.

### Gene expression profiling

We performed gene expression analysis using the NanoString’s nCounter PanCancer Immuno Profiling Panel. RNA was extracted from the FFPE samples using the RecoverAll Total Nucleic Acid Isolation Kit (Ambion; #AM1975). Quality check (QC) was performed for all RNA samples on an Agilent’s 2100 Bioanalyzer. The amount of RNA used for the Immuno Profiling Panel analysis was adjusted according to RNA’s QC score. Using the NanoString’s nSolver 3.0 software, the data were analyzed following the standard protocol. Prior to normalization, the processed samples were checked for quality using the standard QC protocol. Background subtraction was done following the standard protocol using 8 negative controls. Geometric mean of 40 housekeeping genes was used for normalization. All data, except for volcano plots, were generated by Human PanCancer Immune Profiling Advanced Analysis Software (version 1.0.36). For the statistical analysis, we used pre-defined Benjamini-Yekutieli statistical method (p-value threshold 0.05).

## Supplementary Case Details

### Patient 1

A 29yo was diagnosed with a large ovarian tumor that was completely resected. She received initial treatment with bleomycin/etoposide/cisplatin. She was then disease-free for ∼1.5 years and suffered a recurrence in the abdomen and pelvis that was treated with investigational therapy with disease progression in less than 3 months. She was then treated with local radiation and pembrolizumab. She remains on pembrolizumab and continues to have a sustained partial response for 6 months.

### Patient 2

A 22yo was diagnosed with an ovarian tumor completely resected. She did not receive and initial adjuvant treatment and remained disease-free for one year when a recurrence was found in the abdomen and pelvis. She was treated with etoposide/cisplatin followed by surgical resection and then platinum/taxane therapy followed by abdominal RT. She remained disease-free for 9 months and suffered an upper abdominal recurrence treated with RT. She then went on an investigational vaccine study and recurred 12 months later. She started nivolumab and remains disease-free for more than 1.5 years.

### Patient 3

A 25yo was diagnosed with an ovarian mass that was completely resected. She received initial treatment with doxorubicin / cyclophosphamide /etoposide/cisplatin followed by RT. She was then disease-free for ∼1 year and suffered a recurrence in the abdomen. This was surgically resected and taxane/platinum adjuvant therapy was given with concurrent RT. She remained disease-free for six months when she suffered a recurrence in the same area that was treated with additional RT followed by nivolumab. She remains disease free for ∼1.5years.

### Patient 4

An 18yo was diagnosed with a large ovarian tumor that was completely resected. She received initial treatment with etoposide/platinum followed by chemoRT to the abdomen. She was then disease-free for ∼3 years and suffered a recurrence in the chest that was treated with PARP inhibition followed by retreatment with etoposide/platinum, an investigational agent, local radiation and then two cycles of nivolumab. Nivolumab was stopped due to an exacerbation of rheumatoid arthritis and she remains off treatment for more than 1.5 years without any evidence of disease progression.

## Supplementary Figure Legends

**Supplementary Figure 1.**
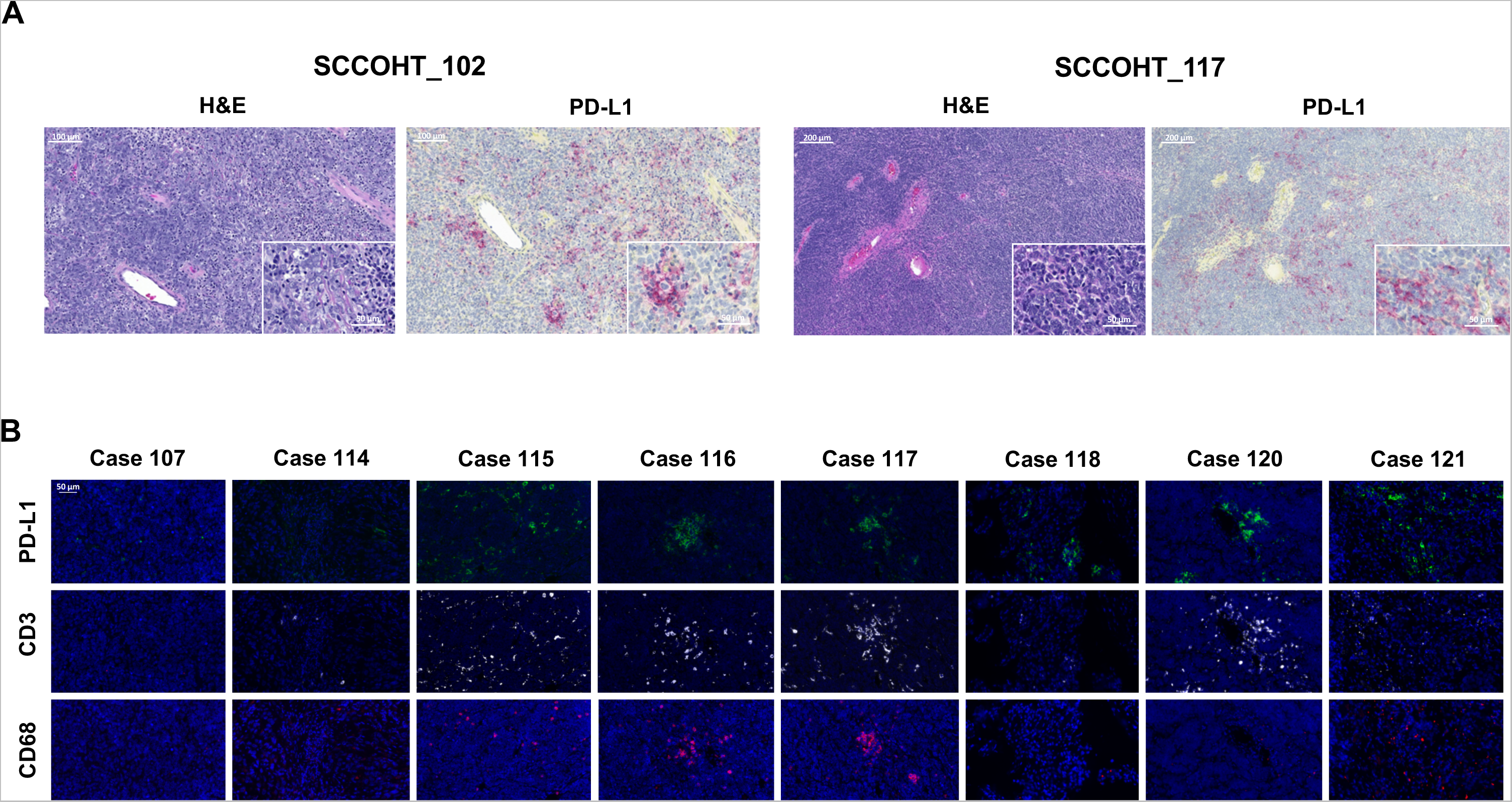
T cell and macrophage infiltration and PD-L1 expression in SCCOHT. **A)** Two representative SCCOHT cases, 102 and 117, showing H&E staining (left) and PD-L1 IHC single stains (right). Inset shows higher power (400X). **B)** IF staining with PD-L1, CD3 and CD68 antibodies for 8 SCCOHT cases.

**Supplementary Figure 2.**
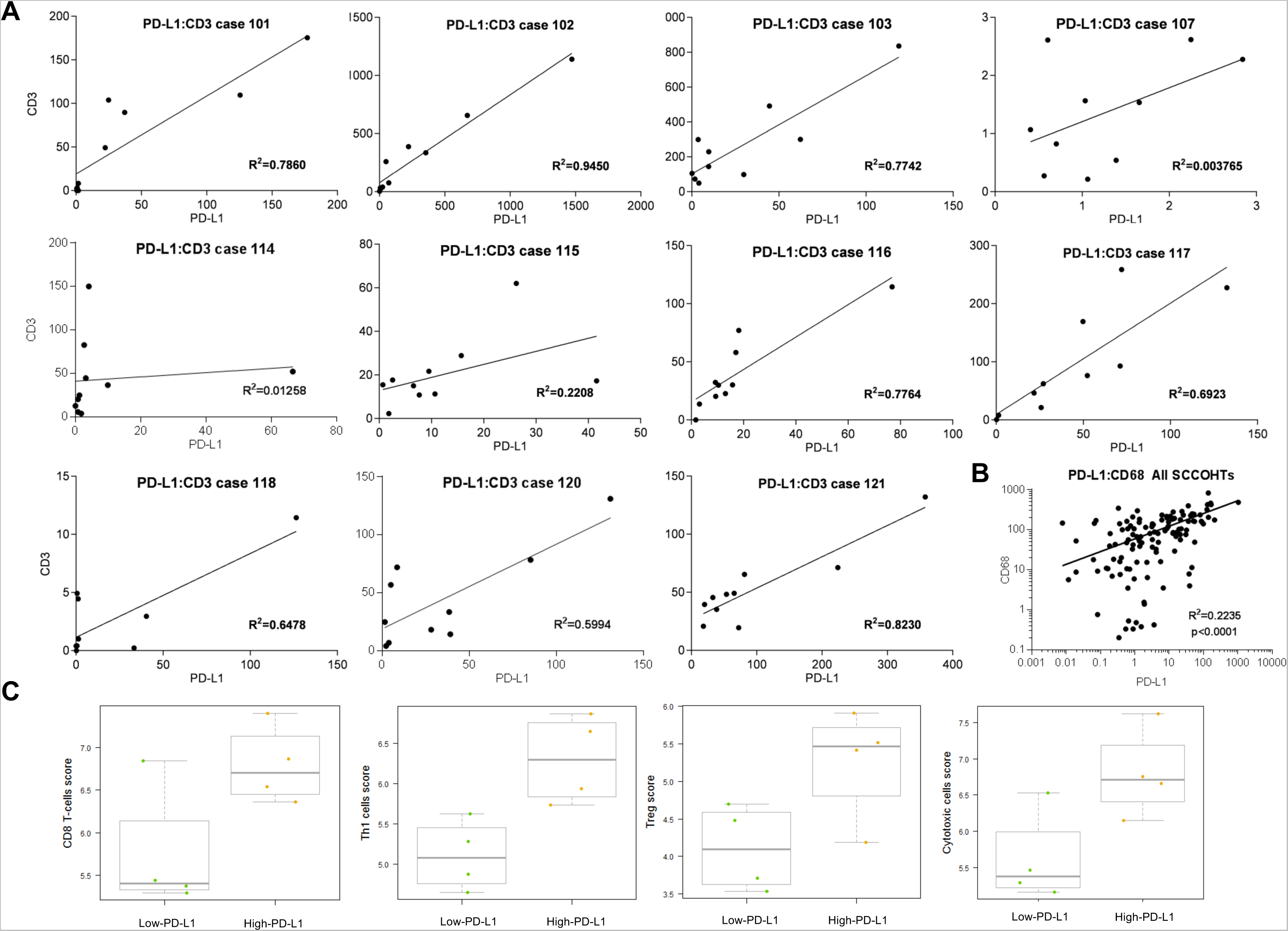
PD-L1 expression correlates with T cell infiltration in SCCOHT. **A)** Correlation of PD-L1 and CD3 expression in SCCOHT cases determined by IF. The y-axes and x-axes represent number of CD3- and PD-L1-positive cells per HPF, respectively. Each dot represents 1 HPF. **B)** Correlation between PD-L1 and CD68 expression in SCCOHT cases. **C)** T-cell subtype scores for High and Low PD-L1 subgroups. Boxplot center lines represent tumor medians, box limits are the interquartile range from 25% and 75%, whiskers represent the extent of tumors out to 1.5 times the inter-quartile range. Dots represent individual cases.

**Supplementary Figure 3.**
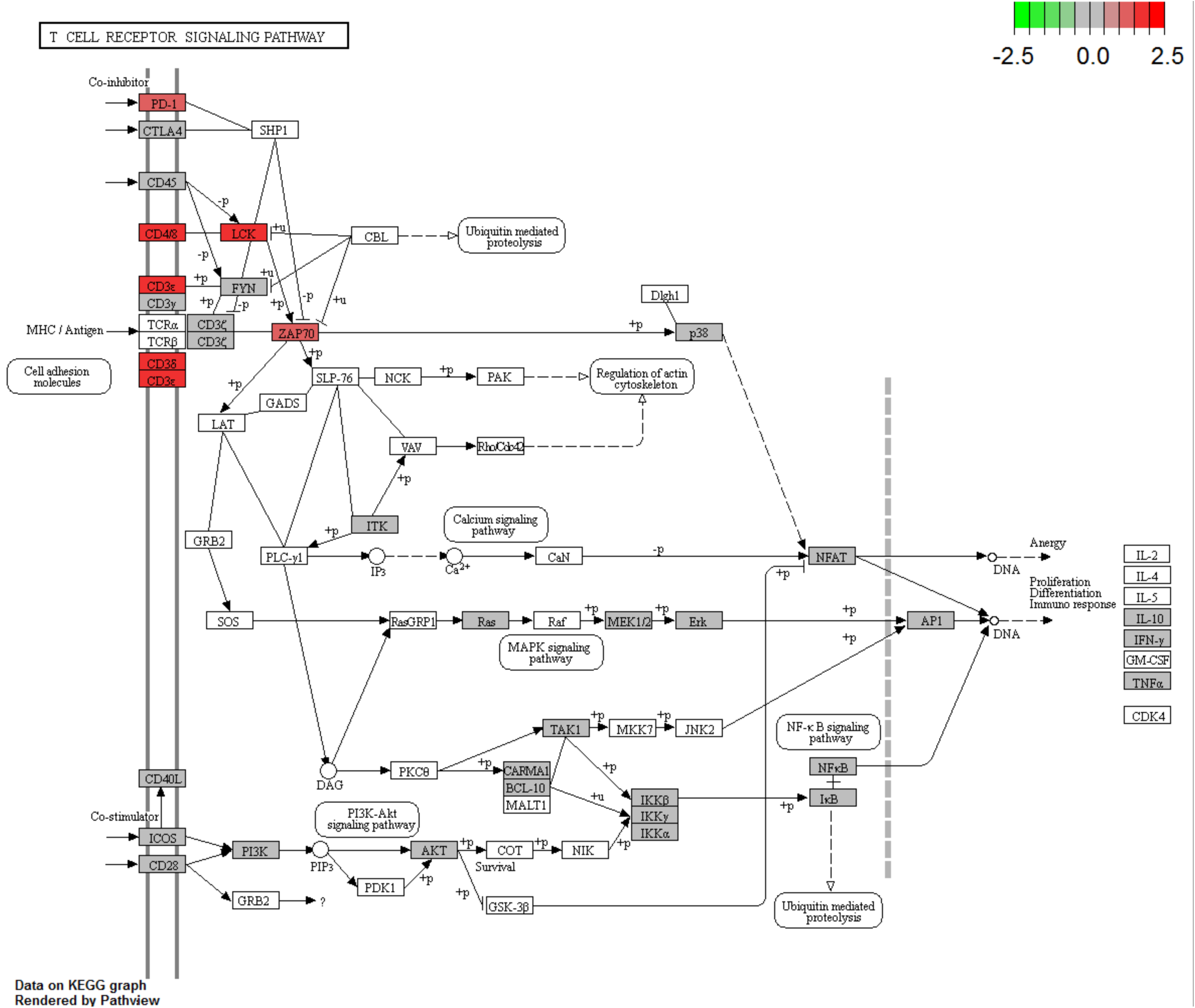
The KEGG graph demonstrates that the T-cell receptor signaling pathway activity is elevated in High- versus Low-PD-L1-expressing SCCOHTs. Red boxes indicate over-expressed genes. Grey boxes indicate genes without differential expression. White boxes indicate genes without expression data.

**Supplementary Table 1.** Summary of *SMARCA4* mutation and protein expression in SCCOHTs.

| Case number | Age at diagnosis (years) | SMARCA4 |  |  |
| --- | --- | --- | --- | --- |
|  |  | Gene mutations |  | IHC |
|  |  | Protein Change | Domain affected |  |
| **101\*** | 40 | p.Q1182_splice | Helicase | Loss |
| **102\*** | 22 | p.K1390_splice |  | Loss |
| **103\*** | 19 | In-frame Deletion | Helicase | Retained |
| **107\*** | 18 | p.E1300_N1303.del |  | Loss |
| **114\*** | 27 | p.X991_splice | SNF2 | Loss |
| **115** | 29 | p.L792P and p.X587_splice | SNF2 | Equivocal |
| **116** | 30 | p.X813_splice | SNF2 | Equivocal |
| **117** | 32 | p.M649Nfs*6 and pE488* |  | Loss |
| **118** | N/A | N/A |  | Loss |
| **120** | 31 | p.X1072_splice | Helicase | Loss |
| **121** | 24 | p.X1318_splice |  | Equivocal |

IHC (immunohistochemistry), SNF2 (sucrose nonfermenting)
\*these data were previously reported [1, 2]

**Supplementary Table 2.** Differentially expressed genes in High- versus Low-PD-L1 expressing SCCOHTs. Cytotoxicity and T-cell Functions gene sets are in bold.

| GENE | Fold change | p-value | Gene sets |
| --- | --- | --- | --- |
| **PRF1** | 3.25 | 0.00076 | **Cytotoxicity**, Pathogen Defense |
| **NCR1** | 3.71 | 0.0032 | Cell Functions, NK Cell Functions |
| **TIGIT** | 2.79 | 0.0058 | **T-Cell Functions** |
| **CLEC4C** | 4.79 | 0.0060 |  |
| **XCR1** | 2.00 | 0.0062 | Chemokines |
| **TNFRSF9** | 2.68 | 0.0065 | TNF Superfamily |
| **CXCL13** | 6.32 | 0.0097 | Chemokines |
| **MEF2C** | -2.51 | 0.012 |  |
| **CXCL9** | 3.03 | 0.015 | Chemokines, Regulation, **T-Cell Functions** |
| **CD8A** | 3.27 | 0.017 | Antigen Processing, Pathogen Defense, **T-Cell Functions** |
| **CXCR3** | 2.85 | 0.018 | Chemokines, **T-Cell Functions** |
| **IL12RB1** | 2.16 | 0.019 | **T-Cell Functions** |
| **CD27** | 4.92 | 0.019 | B-Cell Functions |
| **CD2** | 3.51 | 0.019 | Leukocyte Functions, **T-Cell Functions** |
| **IL6R** | 3.29 | 0.019 | Cytokines |
| **CTSW** | 3.01 | 0.021 | Transporter Functions |
| **GNLY** | 4.56 | 0.025 | **Cytotoxicity**, Pathogen Defense |
| **PDCD1** | 2.30 | 0.025 | Regulation, **T-Cell Functions** |
| **NLRC5** | 3.53 | 0.027 |  |
| **GZMB** | 4.26 | 0.027 | Cell Functions, **Cytotoxicity** |
| **KLRK1** | 2.60 | 0.028 | NK Cell Functions, Regulation |
| **CCL5** | 3.41 | 0.028 | Chemokines, Cytokines |
| **ZAP70** | 2.83 | 0.029 | **T-Cell Functions** |
| **SLAMF7** | 2.60 | 0.030 |  |
| **KLRD1** | 2.79 | 0.032 | NK Cell Functions, Regulation |
| **IDO1** | 3.32 | 0.033 | Cytokines, **T-Cell Functions** |
| **CD38** | 3.43 | 0.033 | B-Cell Functions, Regulation |
| **EWSR1** | 1.12 | 0.033 | Cell Functions |
| **SH2D1A** | 3.10 | 0.036 |  |
| **GZMH** | 2.51 | 0.038 | Cell Functions, **Cytotoxicity** |
| **CD9** | -2.08 | 0.039 |  |
| **BATF** | 2.53 | 0.039 | Cell Functions |
| **CD3D** | 3.53 | 0.040 | Regulation, **T-Cell Functions** |
| **LCK** | 3.29 | 0.041 | Regulation, **T-Cell Functions** |
| **CD5** | 4.26 | 0.044 | Regulation, **T-Cell Functions** |
| **CD3E** | 3.12 | 0.045 | **T-Cell Functions** |
| ***** |  |  |  |
| **CD274** | 1.89 | 0.092 | **T-Cell Functions** |

